# Biofilm Forming Capacity, Sanitizer Tolerance, and Genetic Characterization of Persistent and Non-Persistent *Listeria monocytogenes* from Artisanal Cheese Processing Environments

**DOI:** 10.1101/2025.01.03.631244

**Authors:** Eurydice Aboagye, Emily Forauer, Aislinn Gilmour, Hannah B. Blackwell, Lara Cushman, Calleigh Herren, Sophia Denaro, Lily Felten, McKenzie Ballard, Andrea Etter

## Abstract

*Listeria monocytogenes* is known to colonize food production environments and cross-contaminate finished foods. We investigated 30 *L. monocytogenes* collected from artisan cheese production facilities in Vermont from 2006-2008 for sanitizer tolerance, biofilm formation capacity, biofilm architecture, and tolerance to sanitizers of mature biofilms. Sixteen of these isolates represented a putatively persistent ribotype (DUP-1042B) found in one facility over two years. Isolates of the putatively persistent ribotype all aligned into ST191 and were 0-6 SNPs different, confirming they represented a persistent strain. We found no significant differences in sanitizer tolerance or crystal assay-based attachment capacity between persistent and non-persistent strains. However, using scanning electron microscopy, we found that isolates FML-10 and FML-19 formed substantially denser biofilms after 10 days on stainless steel. Ten-day old biofilms were highly resistant to sanitizers; neither quaternary ammonium nor sodium hypochlorite-based sanitizers achieved an EPA-recommended 6-log reduction. More EPS was found in low-nutrient biofilm conditions; thus, non-food contact surfaces in cheese environments may induce formation of biofilms with high sanitizer tolerance. Our results highlight the importance of regular environmental testing and strain typing for rapid detection of *L. monocytogenes* colonization attempts while they can still be removed without major renovations or equipment replacement.

**Highlights:**

- Isolates from persistent ribotype DUP-1042B/ST191 were within 6 SNPs of each other
- Two isolates from ST191 made dense biofilms in nutrient rich conditions
- More EPS was produced in nutrient-poor conditions
- Mature biofilms of all isolates were highly resistant to QAC and SH sanitizers

**Importance:** This study identifies strategies used by a set of persistently colonizing *L. monocytogenes* isolated from an artisanal cheese producer in Vermont, finding that some persistently colonizing isolates had high biofilm forming capacity, which may have contributed to their persistence.

## Introduction

*Listeria monocytogenes* is a naturally sturdy bacterium that is facultatively anaerobic and capable of growing in broad temperature ranges, from as low as 1.7 ±0.5 °C to as high as 52 °C (1) in some temperature-adapted strains. It is, therefore, ubiquitous in the environment and no stranger to food processing environments (2–7). *L. monocytogenes* is the causative agent of listeriosis, a rare form of bacteremia, meningitis, and encephalitis that is the leading cause of hospitalization from foodborne infection in the United States and is third only to non-typhoidal *Salmonella spp*. and *T. gondii* in terms of mortality (8). When *L. monocytogenes* successfully colonizes and establishes persistence in a food environment, removing it can be extremely challenging and it is more likely to cross-contaminate a finished ready-to-eat product (9–13).

One of the most important pathways through which *L. monocytogenes* establishes persistence is biofilm formation (14–16). Biofilms are complex microbial communities that are attached to a surface and embedded in a protective extracellular matrix (8). The cells in this community develop an altered growth rate and transcriptional pattern, leading to a uniquely stress-tolerant phenotype (17). *L. monocytogenes* biofilms are known to have an increased ability to resist antimicrobial agents and sanitizers and an increased tolerance for high temperatures, low pH, low water activity, UV rays and almost every other avenue for their control in food processing environments (14, 17). Perhaps because of this stress tolerance, biofilm-associated bacteria have been tied to numerous outbreaks. Cherifi et al. found that persistent pulsotypes of *L. monocytogenes* in slaughterhouses in Canada were more likely to be detected in food and human clinical cases than transient strains (18). A six year-long listeriosis outbreak involving Denmark, Estonia, Finland, France and Sweden, was also traced back to a strain *of L. monocytogenes* which was found to be persistent in the premises of a fish smoking facility (19). In the United States, an outbreak associated with ice cream infected at very low levels with *L. monocytogenes* is likely to have been caused by biofilms in the processing environment; in this case, >99% of 2,320 samples of ice cream from one processing facility tested positive for *L. monocytogenes* (9, 10, 20). There is, therefore, ample evidence of the risk of persistent strains of *L. monocytogenes* contaminating finished products to cause outbreaks.

*L. monocytogenes* persistence can be disproportionately devasting for artisanal dairy processors who, unlike their industrial-scale counterparts, do not have the legal and media power to recover from a mass product recall or the loss of consumer confidence (13, 21–23). Thus, outbreaks of listeriosis from these small operations hold not only significant public health concerns, but also the potential loss of important livelihoods and cultural identities tied to artisanal cheese making (13, 21, 24). There is, however, a dearth of information on the phenotypic and genotypic characteristics of persistent *L. monocytogenes* in the context of small-scale dairying operations.

D’amico and Donelly (25), conducted environmental sampling of nine farmstead cheesemaking operations, all of whom produced raw milk cheese and obtained raw milk from their own animals on the same farm on which they were raised. The first sampling event, spanning three months, was conducted in 2006 (25) and found that one facility had a highly prevalent and recurring ribotype (ECORI 210-506-S-5/DUP-1042B). In 2008, most of those same facilities were again sampled in a separate follow-up study (26). D’amico et al. identified a few resident *Listeria spp*. strains which were recovered from multiple sites across all sampling events (26). Most significantly, they identified a specific ribotype of *L. monocytogenes* (DUP-1042B) that was repeatedly isolated from the same facility (facility F in 2006 and Facility A in 2008) and at the same sampling sites (predominantly floors and drains) in both the first sampling event, and then two years later (25, 26). This strain persisted despite improved cleaning and sanitation regimens implemented within the facility and measures to decrease cross-contamination events between the farm and the processing area (26).

We inherited Cathy Donnelly’s extensive *L. monocytogenes* collection, including strains from her 2006 and 2008 studies of artisan dairy environments (25, 26), and we hypothesized that the DUP-1042B ribotype putatively persistent isolates were persisting through enhanced biofilm formation ability(4, 5, 14). We therefore investigated the biofilm-forming capacity as well as the architecture of the resulting biofilms of the DUP-1042B ribotypes from the two D’amico and Donnelly studies (25, 26) grown under nutrient rich (1X) and nutrient poor (1/20X) conditions, and conducted whole genome sequencing to determine the genetic relationships and quaternary efflux pump carriage of these isolates.

## Materials and Methods

### Storage, Recovery and Preparation of Working Stock of Isolates

Frozen stocks of 30 *Listeria monocytogenes* isolates, stored in 25% glycerol and Trypticase Soy Broth (TSB, BD, Franklin Lakes, NJ) at −80°C, were streaked onto Brain Heart Infusion (BHI, Difco, Life Technologies, Detroit, MI) agar and incubated at 37°C for 24 hours. An isolated colony was then selected from each to inoculate BHI broth and incubated at 37°C with shaking (200 rpm) for 18-24 hours. The overnight broth culture was added at a 1:1 ratio of culture to 50% glycerol in cryovials for storage at - 80°C as working stock for the duration of these experiments.

#### Microtiter Assay for planktonic *Listeria monocytogenes* Sanitizer Tolerance

Minimum Inhibitory Concentration (MIC) for sanitizers was determined based on methods from Wang et al. (121). Isolates were recovered from the working stock solutions by streaking 5µL onto BHI agar and incubating at 37°C overnight, after which isolated colonies were re-suspended and adjusted to a range of OD_600_ 0.124-0.140, as read with BioTek Epoch microplate spectrophotometer (BioTek, Winooski, VT). The adjusted culture was then diluted 1:100 into 1mL of 2x concentration BHI or 1/10x concentration BHI and further diluted with the addition of sanitizers, resulting in full strength or 1/20^th^ strength BHI to approximate nutrient rich and nutrient poor conditions.

The 96-well plates were thus inoculated resulting in 100µl each of 3200, 1600, 800, 400, 200, and 100ppm sodium hypochlorite (SH; Ecolab XY-12 Disinfectant with Sodium Hypochlorite; Ecolab, Naperville, IL), or 100, 50, 25, 12.5, 6.25, and 3.125ppm quaternary ammonium compound (QAC) sanitizer (J-512 quaternary ammonium sanitizer; Diversey, Fort Mill, SC), and 1X and 1/20X BHI, with negative and positive controls. Plates were read at hour 0 immediately following inoculation to determine baseline OD_600_, and again to measure growth following static incubation for 24 hours at room temperature (22°C). At the 24-hours reading, test wells were compared to negative control OD_600_ measurement, and MIC was determined to be the concentration of sanitizer where no growth occurred. Growth was defined as having a greater value than negative control of respective sanitizer concentration.

#### Standard Crystal Violet Biofilm Assay

Crystal violet attachment assays were performed as previously described (5). Briefly, overnight cultures were diluted in BHI broth and adjusted to between OD_600_ 0.05-0.10 using the BioTek Epoch spectrophotometer (Biotek, Winooski, Vermont). The adjusted culture for each isolate was then added in triplicate to the wells of three 96-well microplates. The negative control was prepared by adding sterile BHI broth to three wells in each plate. The microplates were then incubated at room temperature (22°C) for one, three, or five days. Optical density was to determine crystal violet retention as an approximation of biomass generated.

#### Evaluating Sanitizer Efficacy on Mature *Listeria monocytogenes* Biofilm

##### Preparation of Culture

Frozen stocks of *L. monocytogenes* isolates were streaked onto BHI agar and incubated at 37°C for 24 hours. An isolated typical colony was aseptically selected to inoculate BHI broth. Broth was incubated at 37°C with shaking (200rpm) for 18-24 hours. Liquid culture was added to either full strength media (1X BHI) or reduced strength media (1/20X BHI) and then OD_600_ adjusted between 0.06-0.10, read on the BioTek Epoch microplate spectrophotometer (Biotek, Winooski, Vermont).

##### Preparation of Stainless-Steel Coupons

Stainless-steel coupons (AISI 304, 1.2mm thick, 10mm diameter, machined by UVM Instrumentation and Model Facility) were soaked in acetone for 30 minutes to remove residues from manufacture, then scrubbed with constant agitation in liquid dish detergent for five minutes and rinsed five times with deionized water to remove soap residue, followed by an ethanol rinse before autoclaving at 121°C for 15 minutes to sterilize. Sterile coupons were then aseptically transferred to empty sterile petri dishes prior to inoculation.

##### Formation of Biofilms

Petri dishes containing six coupons each were inoculated with 10mL OD-adjusted liquid culture. Coupons were then incubated statically at room temperature (22°C) for 10 days. To prevent nutrient exhaustion, spent media was removed and BHI broth was replaced on days two, four, six, and eight of incubation. Experiments were repeated three times on separate days.

##### Sanitizer Application and Enumeration

After 10 days, the broth was removed from the petri dish and coupons rinsed three times with 20mL aliquots of phosphatebuffered saline (PBS) to remove unadhered cells. Four rinsed coupons were then transferred using sterile forceps to test tubes containing 1mL of 0, 50, 100, or 200ppm sanitizer. After the manufacturer-recommended contact time of 60 seconds for QAC and 2minutes for SH, Dey-Engley neutralizing broth was added to tubes to halt the action of the sanitizer. Sterile glass beads (2mm, 1 +/-.1g) were then added and vortexed for two minutes to remove adherent cells into suspension. Samples were serially diluted in buffered peptone water, and spot plated onto BHI agar. Plates were incubated for 24 hours at 37°C prior to enumeration.

#### Scanning Electron Microscopy (SEM) Imaging

Two coupons, one treated with 200pm sanitizer (QAC or SH) and the other rinsed with sterile distilled water (control) were prepared for SEM imaging. Biofilm tissues were fixed in Karnovski’s fixative (0.1M cacodylate buffer, pH 7.2) for 60 minutes at 4°C. Coupons were then rinsed three times for five minutes at 4°C in 0.1M cacodylate buffer, PH 7.2. Biofilms were then fixed with 1% osmium tetraoxide in 0.1M cacodylate buffer, pH 7.2 for one hour at 4°C. This was followed by another three-fold rise for five minutes at 4°C in 0.1M cacodylate buffer. The coupons in six well plates were then wrapped in parafilm and stored in a refrigerator. The next day, the coupons were rinsed once for five minutes in 0.05M cacodylate buffer followed by a three-fold rinse for five minutes in double-deonized water (ddH_2_0). Rinsed coupons were then incubated with 0.5% uranyl acetate in ddH_2_0 for one hour at room (22°C) temperature in darkness (covered with a cardboard box). The samples were then serially dehydrated in ethanol as follows: 35% ethanol, 50% ethanol, 70% ethanol, 85% ethanol, 95% ethanol for 10 minutes each, and finally 100% ethanol three times for five minutes.

The dehydrated samples were then critical point dried with liquid CO2 and mounted on an aluminum specimen holder with conductive paint and dried overnight, before sputter coating with AU/Pd and examining in the Jeol JSM-6060 SEM (Peabody, MA) at the UVM microscopy core.

#### Genomic Analysis of Potentially Persistent *L. monocytogenes* from Farmstead Cheese Production Environments

##### Nanopore sequencing

DNA was extracted from overnight cultures of 30 *L. monocytogenes* using the phenol:chloroform extraction protocol. The extracted DNA was prepared for sequencing on the Oxford Minion sequencer (Oxford, United Kingdom), using the ligation kit and prepared for sequencing with standard Oxford nanopore protocols. Post-sequencing, genomes were trimmed for quality and assembled *de novo* using the long-read genome assembler Flye. Genome assembly quality was then assessed with BUSCO, polished with Medaka and then run through BUSCO again to reassess assemblies quality post-polishing.

##### Illumina sequencing

The Vermont Department of Health performed DNA extraction library preparation with Nextera DNA Flex and generated paired-end reads with an Illumina MiSeq instrument in 2022 (Illumina, Inc., San Diego, CA). These data were submitted through Pulsenet to the National Center for Biotechnology Information (NCBI) website (accessions listed in **Table S1**). The paired-end Illumina reads were downloaded with SRA-toolkit v.2.9.6 with the fastq-dump (split-reads option). The sequence quality was analyzed with Fast-QC v.0.11.3 to determine coverage using the reported read length, number of reads, and *L. monocytogenes* reference genome (NC_003210.1) size.

Draft genomes were assembled with SPAdes v.3.15.5 (27). All Center for Genomic Epidemiology (CGE) tools accepted the unmodified Illumina paired-end reads as input files; each program converted the raw reads into *de novo* draft genomes using Velvet and VelvetOptimizer (28, 29).

**Hybrid assembly**: short-read and long-read genomes were combined using Unicycler v.0.5.0 (30). All isolates FML-007 assembled into hybrid genomes; FML-007 failed at the contig assembly stage.

###### *In Silico* Genotyping

Multi-locus sequence typing (MLST) was performed with MLST v.2.0.9 on both Illumina fastq files and hybrid assembly files (31). The MLST classification was used to determine the lineage with schemes available through the Institut Pasteur (32). Typing of extra-chromosomal features was performed with PlasmidFinder v.2.1 (33). Antimicrobial and stress resistant genotypes were detected with the antimicrobial resistance (AMR) surveillance feature on NCBI and ResFinder (34). These searches were not specific to *L. monocytogenes* gene databases, so ubiquitous genes in the *L. monocytogenes* genome that confer intrinsic resistance to antimicrobials (ex. *fosX*) were excluded from the reported results (35). *L. monocytogenes* virulence genes of interest *inlA, inlB,* and *prfA*, were identified with VirulenceFinder v.2.0.3 (36). The presence or absence of *inlB* was noted directly from the output hits. Nucleotide mutations in *prfA* were detected with a BLAST search with the reference gene; isolates with base mutations were tested for amino acid changes with the Expasy translate tool (37). The Expasy translate tool was also used to detect premature stop codons in *inlA*, which were then characterized based on position in the gene (37, 38). We also compared our genotyping results with the predictions generated by the new tool ListPredv.0.2.0 (39) for virulence and sanitizer tolerance.

###### Single Nucleotide Polymorphism (SNP) Analysis

All 16 ST191 isolates were submitted to MINTyper v.1.0 for SNP typing and extraction of high-quality SNPs (40). MINTyper identified *L. monocytogenes* H34 as the best reference for these 16 isolates (40, 41). We also submitted all 30 hybrid genomes to the Danish Technical University’s CSI Phylogeny tool (v1.4), with *L. monocytogenes* H34 (assembly ASM210127v1) used as the reference genome (42). Dendograms were created from Newick files using FigTree (v.1.4.4), available from https://github.com/rambaut/figtree/releases/tag/v1.4.4.

###### Stress survival islet 1 (SSI-1) detection

A BLAST search for highly similar sequences (megablast) was used to determine the presence of SSI-1 with the “align two or more sequences” option. Gene sequences for lmo044-lmo0448 from the *L. monocytogenes* reference genome EDG_e (NC_003210.1) were used as a query against the subject contigs generated from the SPAdes pipeline (43, 44). SSI-1 positive isolates had all genes present with at least 99% identity for each gene.

#### Statistical Analyses

To determine variations in MIC between isolates and between nutrient concentrations, a one-way Analysis of Variance (ANOVA), after satisfying conditions of homogeneity (Levine’s test), and normality (QQ plots), was used. Analysis of covariance (ANCOVA) was used to determine how the mean optical density (a measure of biofilm forming capacity) varied between isolates, with nutrient concentration as a covariate. Finally, to evaluate the efficacy of sanitizers on mature biofilms grown under nutrient rich and poor conditions, the primary outcome variable, log reduction, was determined by subtracting the log_10_CFU/coupon after sanitizer treatment, from the log_10_CFU of the untreated (control) coupon. A factorial ANOVA was then used to determine the effects of sanitizer type, sanitizer concentration and nutrient concentration, as well as the interactions between these variables on log reduction. A Student t-test was additionally carried out to compare the efficacy of the two sanitizers. All statistical analyses were carried out using RStudio 2023.12.1+402 "Ocean Storm" Release for windows Mozilla/5.0 (Windows NT 10.0; Win64; x64) AppleWebKit/537.36 (KHTML, like Gecko) RStudio/2023.12.1+402 Chrome/116.0.5845.190 Electron/26.2.4 Safari/537.36.

## Results

### Whole Genome Sequence of *L. monocytogenes* Isolates from Artisanal Cheese Processing Environment

The 30 *L. monocytogenes* isolates **(Table 1)** clustered into five SNP groups **(Figure 1)**. Isolates from lineage II aligned into four SNP groups. with only one SNP group from lineage I—the ST191 SNP group. ribogroup, ribotype and MLST type largely concurred, but did not fully align. All ECORI 210-506-S-5/DUP-1042/1042B isolates typed as ST191; however, isolate FML-005 (ECORI 210-507-S-6/DUP-1006/1006) also typed as ST191. Conversely, isolate FML-009 (ECORI 210-506-S-1/DUP-19171/1042B) typed as ST361 rather than ST191. Isolates from other ribotypes did not align into MLST types as consistently. All three isolates from ribogroup ECORI 210-511-S-7/18645/1045A typed into ST 222 but isolates from ECORI 210-506-S-1 and ECORI 210-506-S-4 both typed into ST29, and four ECORI 2010-506-S-1 isolates typed into ST361. We used MINTyper to further analyze the 16 ST191 isolates. These isolates ranged from 0-6 SNPs different, indicating they are all the same strain **(Figure 2)**. The isolates’ SNPs were evaluated using MINTyper (40), which aligned them against strain H34 (NCBI reference sequence number: NZ_CP020774.1), a neonatal sepsis strain from Uruguay (41).

**Figure 1:**
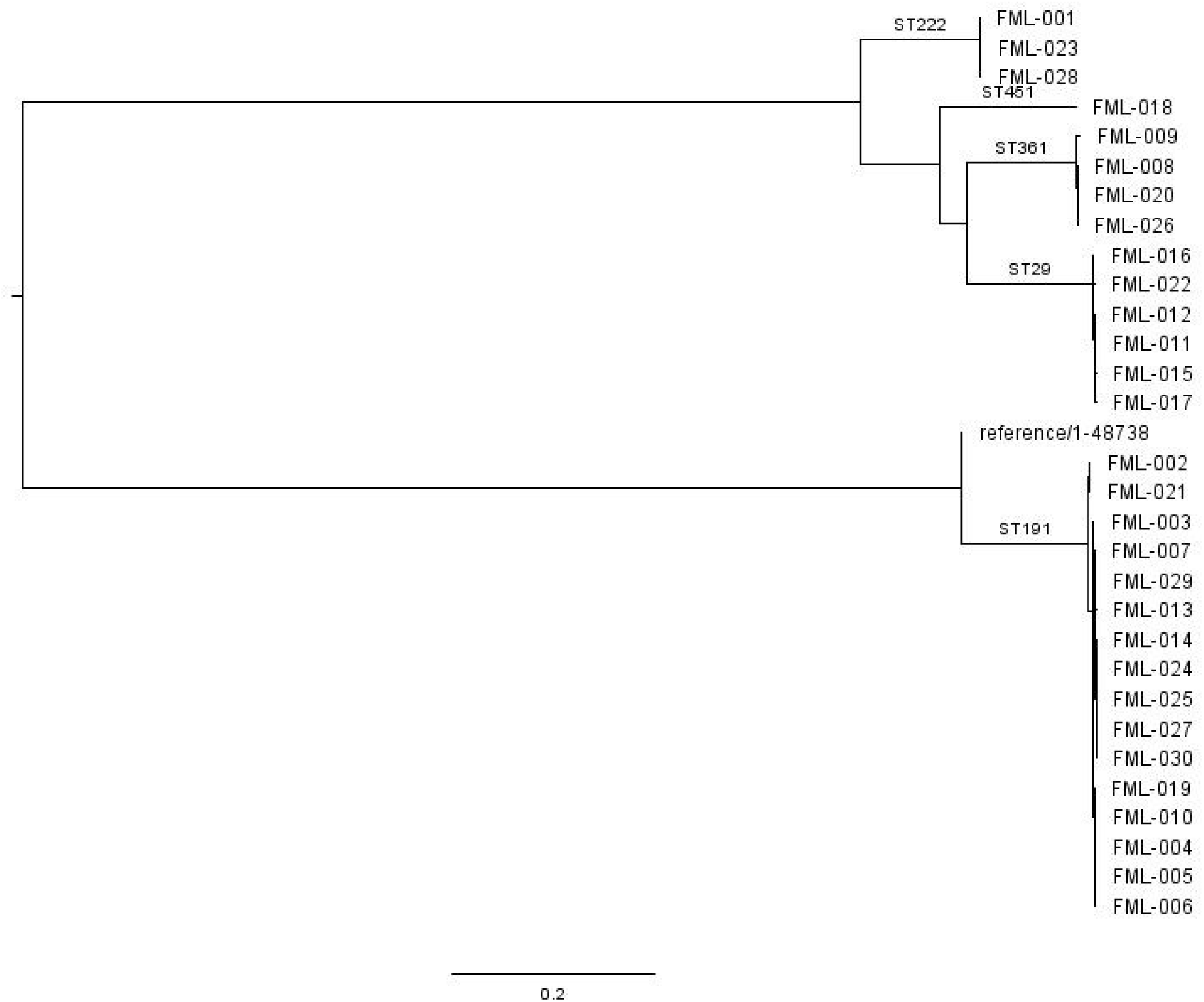
SNP tree of 30 *L. monocytogenes* isolates from this study. Created using CSIPhylogeny (42) with H34 (NZ_CP020774.1) as a reference.

**Figure 2:**
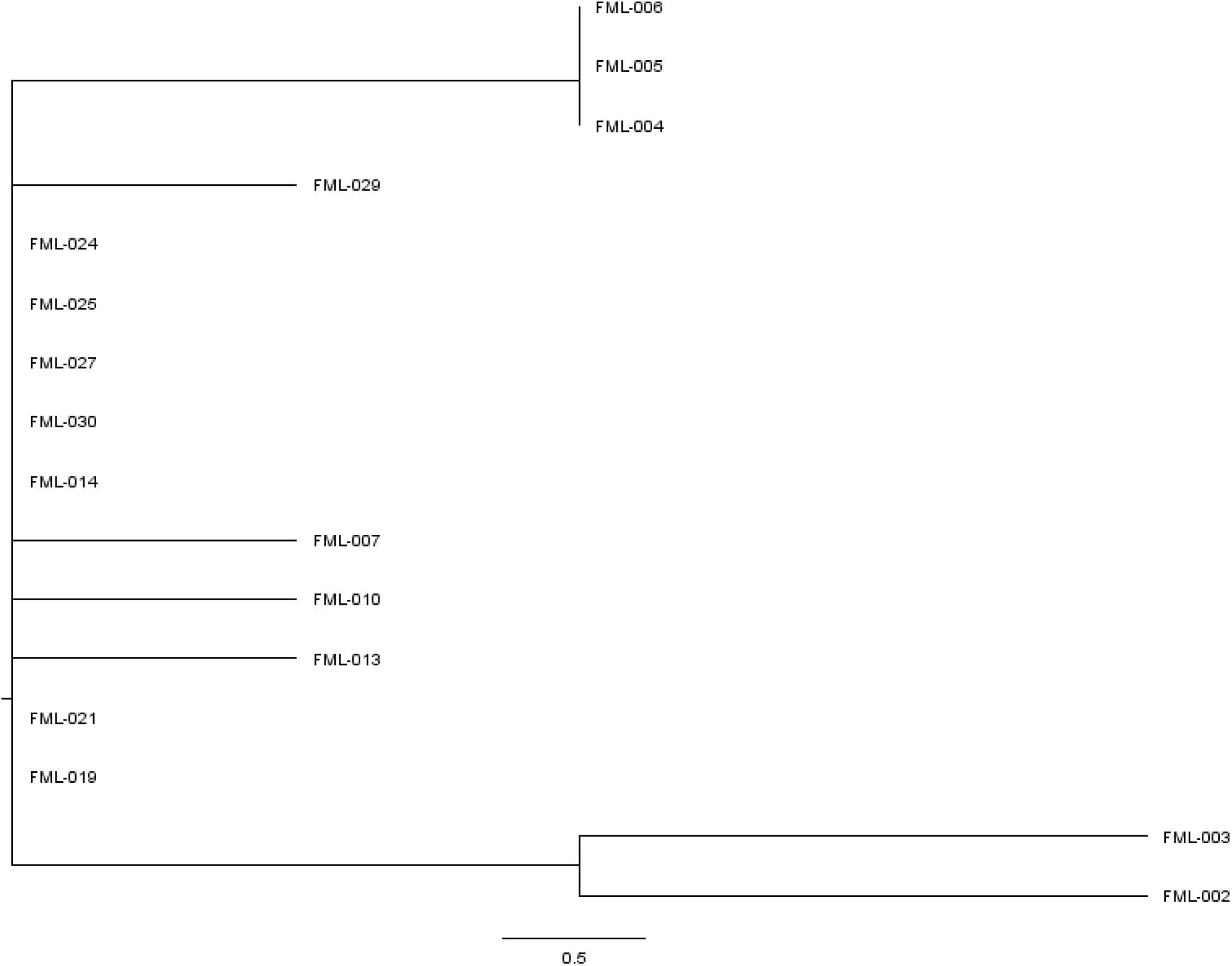
SNP-tree of ST191 isolates. Created using MINTyper (40) with H34 (NZ_CP020774.1) as a reference.

**Table 1:** Minimum Inhibitory Concentrations of Environmental Isolates of *Listeria monocytogenes* in Quaternary Ammonium Compounds (QACs) and in Sodium Hypochlorite-Based Sanitizers.

| Isolate | Date Collected | Environmental Source | Pathogen Tracker ID | MLST Type | Lineage<sup>b</sup> | SSI-1<sup>c</sup> | MIC (ppm) ±SD |  |  |  |
| --- | --- | --- | --- | --- | --- | --- | --- | --- | --- | --- |
|  |  |  |  |  |  |  | SH |  | QAC |  |
|  |  |  |  |  |  |  | Nutrient Poor (0.05X) | Nutrient Rich (1X) | Nutrient Poor (0.05X) | Nutrient Rich (1X) |
| 1 | 10/20/08 | Floor (entrance) | - | 222 | II | - | 100±0 | 666±230 | 5.21±1.81 | 7.03±3.93 |
| 2 | 9/17/08 | Floor (under vat) | 1042B | 191 | I | + | 100±0 | 533±230 | 4.69±1.81 | 6.25±0.00 |
| 3 | 9/17/08 | Drain (aging room 2) | 1042B | 191 | I | + | 100±0 | 533±230 | 3.90±1.57 | 6.25±0.00 |
| 4 | 9/17/08 | Drain (aging room 1) | 1042B | 191 | I | + | 100±0 | 933±611 | 4.69±2.21 | 6.25±0.00 |
| 5 | 9/17/08 | Shoes | 1006 | 191 | I | + | 200±173 | 666±230 | 6.25±0.00 | 7.81±3.13 |
| 6 | 9/17/08 | Floor squeegee | 1042B | 191 | I | + | 100±0 | 533±230 | 6.25±0.00 | 6.25±0.00 |
| 7 | 09/25/06 | Drain (aging) | 1042B | 191 | I | + | 100±0 | 533±230 | 9.38±4.42 | 12.5±8.84 |
| 8 | 09/25/06 | Drain (aging) | 1030B | 361 | II | - | 100±0 | 666±230 | 5.48±1.57 | 9.38±3.61 |
| 9 | 10/16/06 | Floor squeegee | 1042B | 361 | II | - | 200±173 | 666±230 | 5.48±1.57 | 6.25±0.00 |
| 10 | 11/06/06 | Cut and wrap table | 1042B | 191 | I | + | 1133±1789 | 666±230 | 6.25±0.00 | 6.25±0.00 |
| 11 | 10/02/06 | Drain (vat) | 1030B | 29 | II | - | 100±0 | 666±230 | 6.25±0.00 | 6.25±0.00 |
| 12 | 10/02/06 | Drain (vat) | 1039E | 29 | II | - | 100±0 | 1466±1514 | 8.33±3.61 | 8.33±3.61 |
| 13 | 10/16/06 | Pooled water | 1042B | 191 | I | + | 100±0 | 666±230 | 5.21±1.80 | 8.33±3.61 |
| 14 | 10/16/06 | Water hose | 1042B | 191 | I | + | 100±0 | 666±230 | 5.21±1.80 | 6.25±0.00 |
| 15 | 10/02/06 | Drain (vat) | 1038A | 29 | II | - | 2266±1616 | 133±57.7 | 5.21±1.80 | 8.33±3.61 |
| 16 | 10/02/06 | Pooled water | 1023B | 29 | II | - | 360±383 | 1933±1137 | 10.94±3.13 | 18.75±7.22 |
| 17 | 10/02/06 | Pooled water | 1039E | 29 | II | - | 200±173 | 233±152 | 7.81±5.41 | 7.03±3.93 |
| 18 | 10/16/06 | Floor (entrance) | 1039C | 451 | II | - | 1233±1709 | 2400±1385 | 6.25±4.42 | 8.59±4.69 |
| 19 | 10/16/06 | Bucket (fill hoops) | 1042B | 191 | I | + | 1466±1514 | 470±310 | 3.52±1.97 | 4.69±1.80 |
| 20 | 10/16/06 | Floor squeegee | 1044A | 361 | II | - | 200±173 | 933±611 | 3.52±1.97 | 4.69±1.80 |
| 21 | 11/06/06 | Hallway floor | 1042B | 191 | I | + | 133.3±57.74 | 622±203 | 3.52±1.97 | 6.25±4.42 |
| 22 | 10/30/06 | Drain (new) | 1039C | 29 | II | - | 100±0 | 333±404 | 5.47±1.56 | 10.94±3.13 |
| 23 | 10/30/06 | Drain (vat) | 1045A | 222<sup>a</sup> | II | - | 600±866 | 666±230 | 4.30±2.34 | 6.25±4.42 |
| 24 | 09/25/06 | Drain (aging) | 1042B | 191 | I | + | 1133±1789 | 200±173 | 4.69±2.21 | 6.25±0.00 |
| 25 | 10/16/06 | Drain (aging #2) | 1042B | 191 | I | + | 1133±1789 | 333±404 | 3.13±0.00 | 6.25±0.00 |
| 26 | 10/16/06 | Floor squeegee | 1039C | 361 | II | - | 1133±1789 | 844±76.97 | 6.25±0.00 | 9.375±4.42 |
| 27 | 11/06/06 | Cut and wrap<br>table | 1042B | 191 | I | + | 100±0 | 655±499 | 6.25±0.00 | 6.25±0.00 |
| 28 | 10/30/06 | Drain (vat) | 1045A | 222<sup>a</sup> | II | - | 100±0 | 844±76.97 | 6.25±0.00 | 6.25±0.00 |
| 29 | 11/06/06 | Hallway floor | 1042B | 191 | I | + | 50±0 | 844±76.97 | 8.33±3.61 | 15.63±10.83 |
| 30 | 11/06/06 | Cut and wrap<br>table | 1042B | 191 | I | + | 100±86.6 | 1644±1348 | 6.25±0.00 | 6.25±0.00 |

<sup>a</sup>Has a novel allele for *cat*
<sup>b</sup>Based on Pasteur Institute datasheet for MLST types (32)
<sup>c</sup>As described by Keeney et al.(83)

No antimicrobial resistance genes were detected, and none of the isolates had arsenic tolerance gene *arsD* or QAC tolerance genes *bcrABC* or *qacG*. All isolates, regardless of QAC MIC, had the *mdrL* QAC tolerance gene (**Table 2)**; however, this gene has been shown to not substantially affect sanitizer tolerance (45). SSI-1 was present in all lineage I isolates, but it was not detected in any of the lineage II isolates. No isolates had *inlB* gene deletions or *inlA* premature stop codons, indicating all should in theory be fully virulent. Mutations in *prfA* were present in 67% (20/30) of isolates, with all 16 lineage I isolates and four lineage II isolates having identical silent nucleotide substitutions. The *prfA* mutations did not alter the amino acid sequence and were not further analyzed.

**Table 2.** Genetic characteristics of cheese production environment-derived isolates by lineage.

|  | Lineage I (ST191)<br>% (n/total) | Lineage II<br>% (n/total) | Total<br>% (n/total) |
| --- | --- | --- | --- |
| **Stress or AMR gene(s)<sup>1</sup>** | 0 (0/16) | 0 (0/14) | 0 (0/30) |
| **SSI-1 present** | 100 (16/16) | 0 (0/14) | 53 (16/30) |
| **Plasmid(s) present** | 0 (0/16) | 0 (0/14) | 0 (0/30) |
| **PMSC in *inlA*** | 0 (0/16) | 0 (0/14) | 0 (0/30) |
| ***prfA* mutation (nucleotide only)** | 100 (16/16) | 29 (4/14) | 67 (20/30) |
| **ListPred predicted virulence:** |  |  |  |
| **High** | (/16) | (/14) | (/30) |
| **Moderate** | (/16) | (/14) | (/30) |

<sup>1</sup>Genes tested for: *arsD*, *bcrABC*, *qacG*

**Table 3:** Predictors of Sanitizer Tolerance in Mature Biofilms of Listeria monocytogenes.

| Effects | Mean Log Reduction ±SE | P-value |
| --- | --- | --- |
| **Sanitizer Type** |  | **2.2e-16** |
| SH | 2.766±0.151 | |
| QAC | 0.891±0.250 | |
| **Sanitizer Concentration\*Sanitizer** |  | **0.00427** |
| Sub-optimal (QAC) | 0.809±0.122<sup>a</sup> | |
| Optimal (QAC) | 1.055±0.2499<sup>a</sup> | |
| Sub-optimal (SH) | 2.565±0.1127<sup>b</sup> | |
| Optimal (SH) | 3.162±0.151<sup>c</sup> | |
| **Nutrient Concentration** |  | **0.2285** |
| 1X | 2.161±0.298 | |
| 1/20X | 2.478±0.236 | |
| **Nutrient Concentration\* Sanitizer(200pm)** |  | **0.0318** |
| 1X (QAC) | 0.545±0.153<sup>a</sup> | |
| 1X (SH) | 3.237±0.268<sup>b</sup> | |
| 1/20X (QAC) | 1.564±0.437<sup>a</sup> | |
| 1/20X (SH) | 3.0863±0.146<sup>b</sup> | |

Within each effect, means with different superscripts are significantly different at $\alpha < 0.05$ . The effects of Nutrient concentration on the model was determined with each sanitizer at 200ppm (manufacturer recommended levels). Suboptimal levels of sanitizer were studied at 50ppm and 100ppm.

We further ran the assembled hybrid genomes of these isolate through the new machine learning-based tool ListPred, to test the tool’s prediction for our isolates’ sanitizer tolerance and virulence (39). ListPred accurately assessed all of our isolates as QAC susceptible, but it predicted that isolates from ST361 had high virulence, while the remainder of the isolates (STs 29, 191, 222, and 451) were classified as moderately virulent (39). As we did not detect any mutations associated with reduced virulence in the ST191, 222, and 29 isolates, this prediction is more likely based on the clinical frequency of these sequence types than any specific mutations.

### Sanitizer Tolerance of Planktonic Cells

At room temperature (22°C), average minimum inhibitory concentrations (MIC) of QACs in nutrient rich conditions ranged from 4.69ppm to 18.75ppm, and from 3.12ppm to 10.94ppm in nutrient poor conditions **(Table 1).** These MICs were significantly lower than the recommended working concentration of 200ppm for food contact surfaces and 400ppm for non-food contact surfaces, suggesting a rather low tolerance of the planktonic *L. monocytogenes* isolates to QAC. Isolates FML-010-FML-016 and FML-029 had significantly higher QAC tolerance than the other isolates in both nutrient rich (1X) and nutrient poor (1/20X) conditions (p > 0.05) **(Figure S1(a)).** Notably, there were no significant differences (p=0.1794) between the average MIC of isolates grown in nutrient rich vs nutrient poor conditions.

MICs for SH ranged between 100-800ppm in nutrient poor conditions. These MICs, however, were significantly lower than the MICs in nutrient rich conditions, which ranged between 400-1600ppm (p=0.004). These values suggest a high tolerance of the planktonic *L. monocytogenes* to SH. There were no statistical differences among individual isolate MICs, however isolate 16 had greater SH tolerance in both nutrient rich and nutrient poor conditions **(Figure S1 (b)).**

### Attachment Capacity of Isolates (Crystal Violet Assay)

Cell density/ numbers within biofilms of *L. monocytogenes* as measured by optical density (OD) did not vary significantly over the 5-day incubation period. Biomass levels at 24 hours did not significantly change on days 3 and 5. Importantly, isolates grown in nutrient poor conditions (1/20X BHI) had significantly higher biomass (p = 4.386e-07) compared to isolates grown under nutrient rich conditions (1X BHI) (**Figure 3**). Between isolates, there were insignificant variations in cell populations within biofilms formed except in the case of isolate 10. This isolate importantly belongs to the persistent 1042B ribogroup from a single artisan cheese facility and had a mean OD of 0.1876±0.1787 which was significantly higher (p = 0.00038) compared to the rest of the isolates (**Figure 4**).

**Figure 3:**
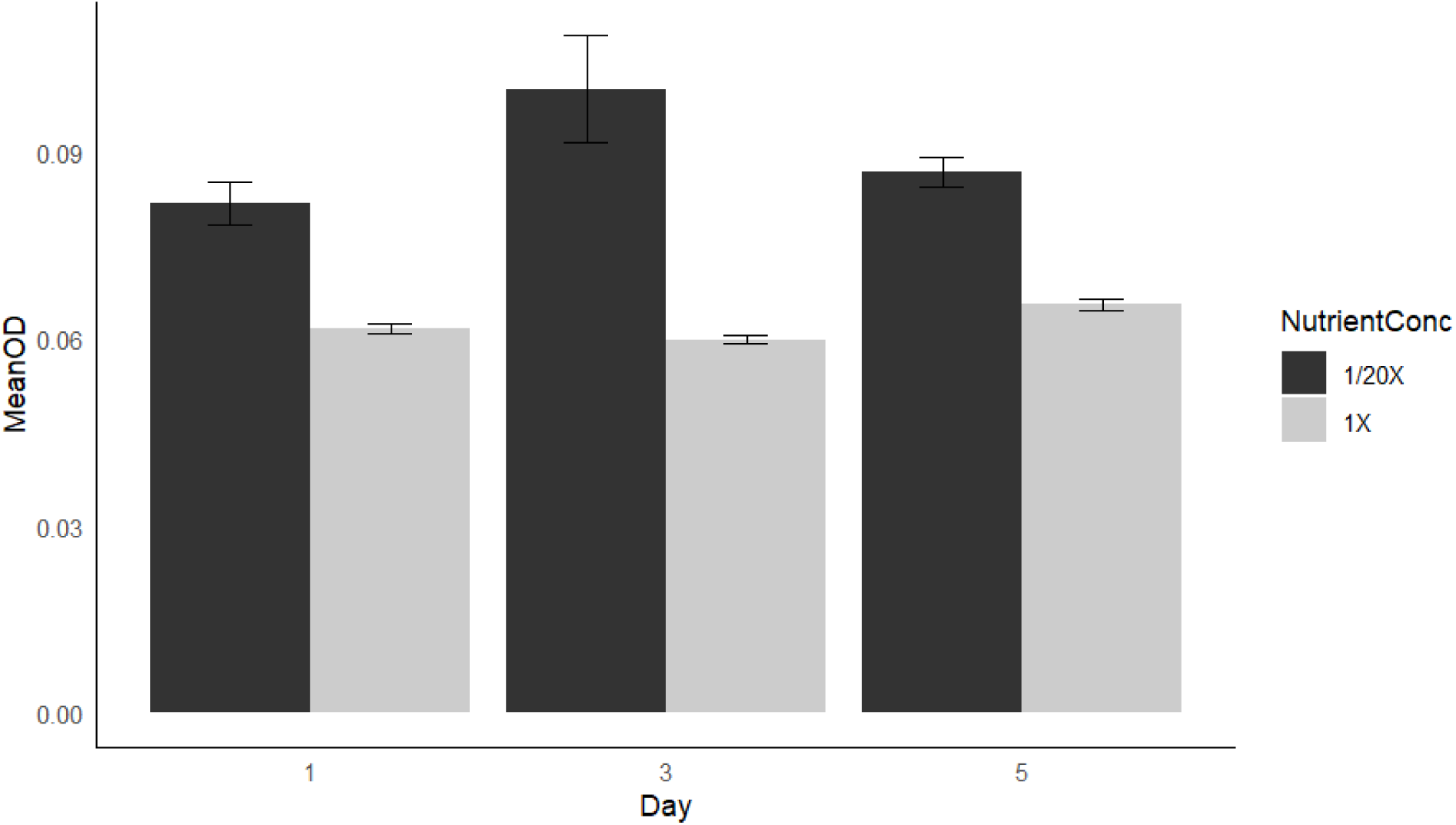
Biofilm forming capacity of environmental *L. monocytogenes* in nutrient rich and nutrient poor conditions at 22°C.

**Figure 4:**
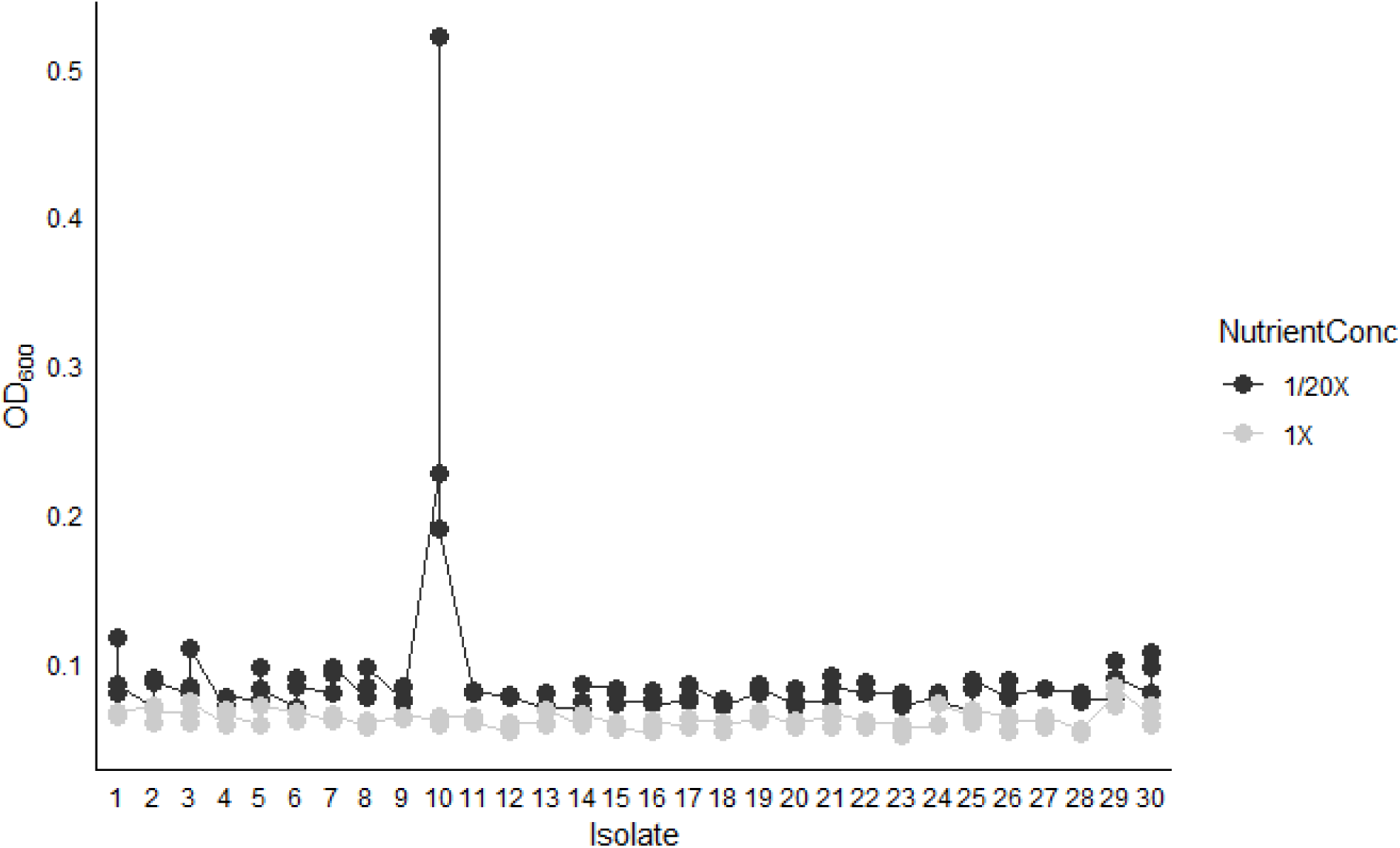
Biofilm forming capacity of *L. monocytogenes* isolates as determined by biomass in nutrient rich and nutrient poor conditions at 22*°*C.

### Mature (10-day) *Listeria monocytogenes* Biofilms on Stainless Steel

We selected six of the 30 isolates to further analyze their ability to form mature biofilms on stainless steel, as this is a common material in the dairy and artisan cheesemaking environment. Isolates were chosen to represent a mix of potentially persistent (DUP-1042) and non-persistent subtypes. We assessed both nutrient-rich (1X BHI) and nutrient poor (1/20X BHI) conditions at room ^i^temperature (22°C). The mean cell population within nutrient rich mature biofilms of the six selected isolates was 7.704 log_10_ CFU/coupon which was significantly greater (p-value < 2.2e-16) than the 6.64797 log_10_CFU/coupon cell population in nutrient poor mature biofilms. Among isolates, however, there was no significant difference between cell populations within the biofilms **(Figure 5).** We also did not find any significant difference between cell counts of the persistent ST191 strain and the transient strains.

**Figure 5:**
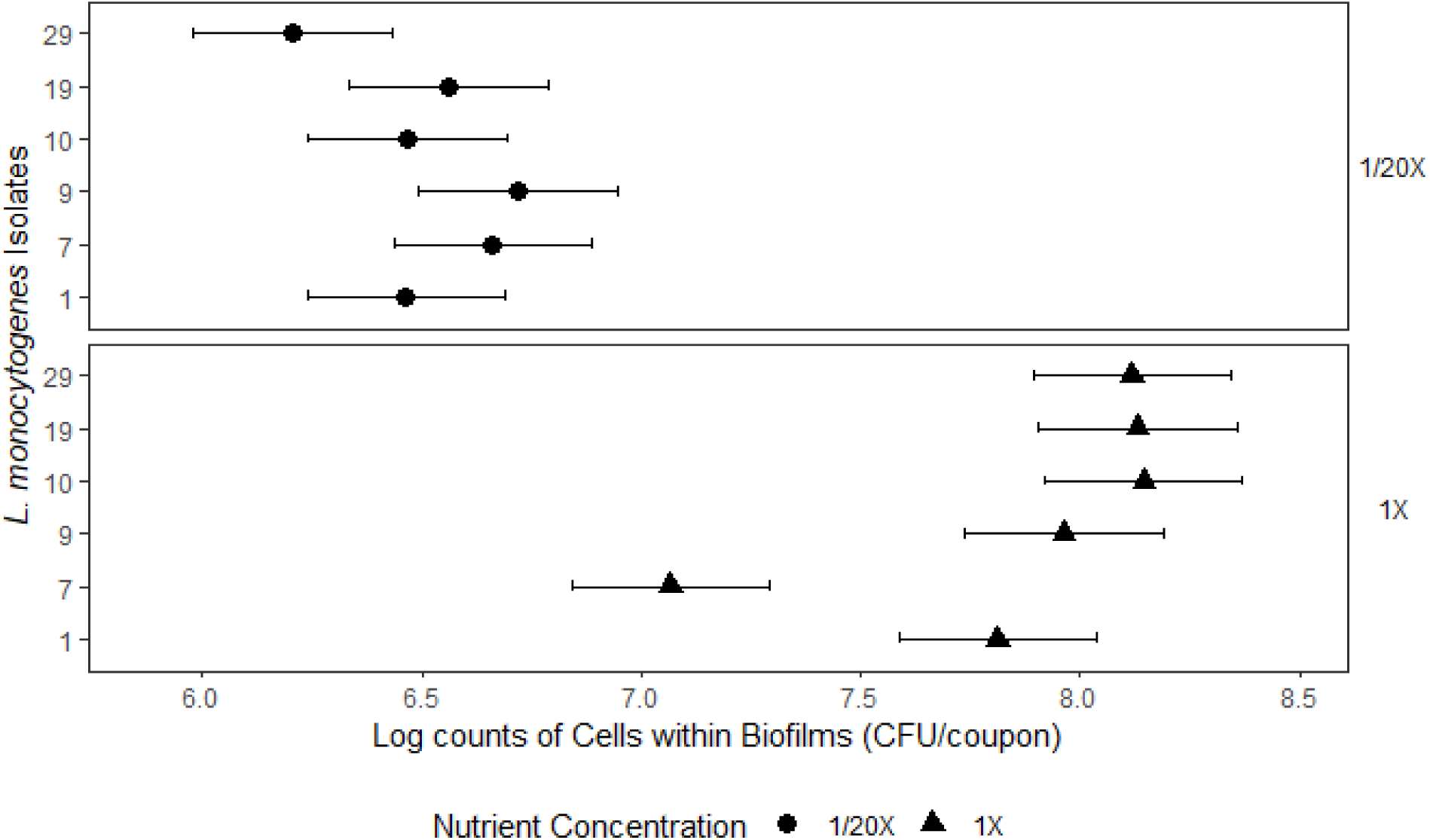
cell counts of mature (10 day) *L. monocytogenes* biofilms from six strains on stainless steel.

### Biofilm architecture

We used Scanning Electron Microscopy (SEM) to investigate the biofilm architecture of mature biofilms formed under nutrient rich and nutrient poor conditions at room temperature (22°C), using the same subset of six isolates. There was substantial variation in biofilm architecture under the two different nutrient availability conditions. Nutrient rich biofilms, typically showed dense consistent single-cell coverage with more frequent 3D aggregates **(Figure 6a)**. Under nutrient poor conditions, however, mature biofilms of *L. monocytogenes* on stainless steel coupons were composed of infrequent cell aggregates interspersed with single cells wedged inside micro-fissures on the surface of the coupon **(Figure 6b)**. Isolates FML-009, FML-010 and FML-019, two of which were of the DUP-1042B ribotypes (10, 19), and isolated from the floor squeegee, cutting table and cheese fill hoops, respectively, had especially dense coverage under nutrient rich conditions. They also had substantial 3D development within surface micro blemishes on the surface of the coupon. In nutrient poor conditions, these isolates also showed greater exopolysaccharide development than the remaining isolates.

**Figure 6:**
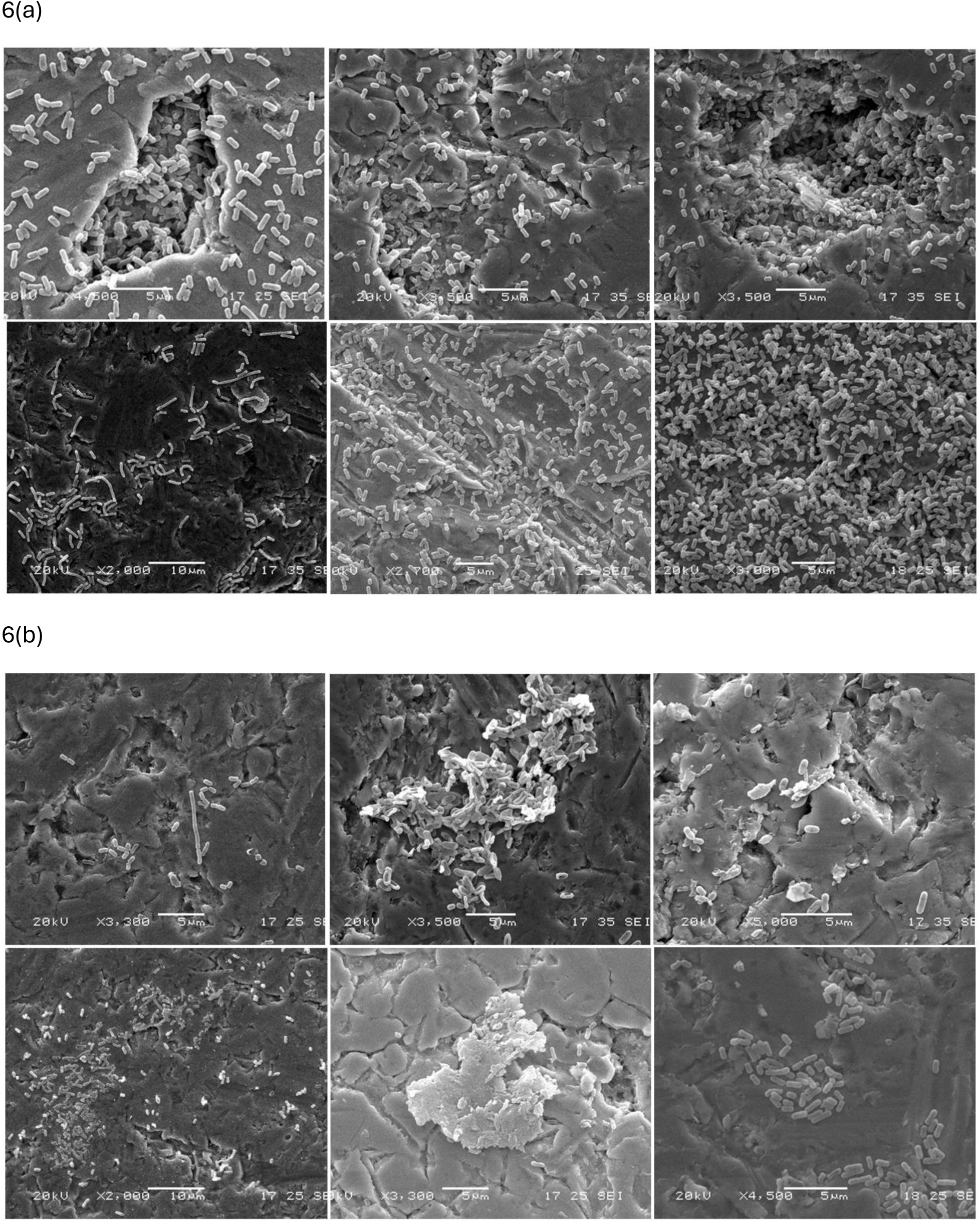
SEM Images of 10-Day *L. monocytogenes* Biofilms (Top row: Isolates 1, 9 and 10; Bottom Row: ST 191 Isolates 7, 19, 29) (a) 1X BHI and (b) 1/20X BHI at 22°C

### Sanitizer Tolerance of Mature *Listeria monocytogenes* Biofilms

Sanitizer tolerance, as determined by the log reduction of cell populations within biofilms after sanitizer application, was significantly affected by sanitizer type, sanitizer concentration and the interaction between nutrient concentration in growth medium and sanitizer used. **Table 2** summarizes the results of a factorial ANOVA model that examined the effects of sanitizer type, sanitizer concentration, nutrient concentration, and their interactions on log reduction of cells simultaneously.

Although both sanitizers were unable to achieve a 6-log reduction, SH was significantly more effective than QACs against mature biofilms, achieving on average, more than 2 Log reductions of cell numbers compared to QACs. Even at sub-optimal levels (50 and 100ppm), SH had a significantly stronger effect against the mature biofilms compared to QAC at manufacturer recommended levels (200ppm). SH was, however, most efficacious at 200ppm.

A significant interaction was also observed between the nutrient concentration of the biofilm growth medium and the sanitizer used. Biofilms grown under nutrient poor conditions were significantly more susceptible to QAC than those grown under nutrient rich conditions. The same relationship was not observed for SH, which was equally effective against nutrient poor biofilms as they were against nutrient rich biofilms.

## Discussion

We sought to investigate (i) whether the DUP-1042B ribotype previously repeatedly re-isolated in a single Vermont artisanal cheesemaking environment represented a persistent strain by using more sensitive strain typing techniques, (ii) whether the repeated re-isolation of this ribotype could be attributed to an inherent genetic or inherent phenotypic trait that influenced biofilm formation and subsequent sanitizer tolerance and (iii) the biofilm and sanitizer-tolerance capacities of both the DUP-1042B and non-DUP-1042B isolates from Vermont artisan cheesemaking environments.

WGS-based MLST typing (31) of isolates was largely similar to the previously performed ribotyping (25, 26); however, it was not identical. All the DUP-1042 isolates in our dataset, which came from a single facility (25, 26), belonged to ST191 and showed 0-6 SNPs difference from each other; however, one additional isolate from DUP-1006 also typed as ST-191 and grouped with two other DUP-1042 isolates (0 SNPs difference). Previous research by Borucki et al., also determined that six isolates of the DUP-1042 ribotype, grouped into two different MLST types and three different serovars (46). Thresholds from ≤6 high-quality SNPs to up to 22 SNPs have been proposed for identifying the same strain/persistence via SNP analysis (47–49); given this, the ST191 isolates can be confidently identified as a persistent strain within the facility they came from. It is also not surprising that there were minimal changes over a two-year period, as *L. monocytogenes* strains have been found to persist in food processing environments for up to 12 years, with minimal genetic changes (48–51).

ST191 is a rare type in the Pasteur Institute’s database of *L. monocytogenes* strains (32, 52), with only nine isolates mapping to that type. Of these nine, two were clinical isolates from blood, two were food isolates, two were isolates from soil or compost, two were from animals, and one was from the production environment (53). Additionally, only a handful of published studies exist that include isolates from ST191. In these studies, ST191 were rare, but found among samples collected from bulk tank milk, milk filters, and milk processing equipment (54); from ruminant listeriosis cases in the US (55, 56) and Europe (57); in the meat processing environment in Poland (58); and in trout meat from French fisheries (59). Consequently, the ST191 isolates in our study may well have originated in an infected animal in the farmstead cheese producer’s dairy herd. Whether from there it was introduced to the cheesemaking facility via *L. monocytogenes* in the milk (60) or was brought in on contaminated footwear or equipment is impossible to know; however, given the widespread nature and persistence of the strain in the cheese production facility (25, 26), the company’s sanitation protocols were clearly inadequate to eradicate it once introduced.

Planktonic forms of all 30 environmental Listeria monocytogenes isolates were readily destroyed at concentrations of QAC well below manufacturer recommendations. In contrast, sessile forms showed significant tolerance to sanitizers at manufacturer-recommended levels of use.

Additionally, QACs were significantly more effective against planktonic cells than SH. None of the isolates had significant phenotypic resistance to QACs in planktonic cultures, as indicated by the fact the highest measured MIC was ∼20ppm. This is not surprising, as none of the isolates contained genetic elements associated with increased resistance to QACs (45). Ivanova et al., recently completed a massive analysis of phenotypic and genotypic concordance for *L. monocytogenes* tolerance to QACs; they found that only *bcrABC, emrC, emrE, or qacH* increased QAC tolerance (45). While all our isolates contained *mdrL,* which was previously reported to increase QAC tolerance (61, 62), this gene has been reported by others to be found in both QAC tolerant and QAC-sensitive isolates (45). Our data also suggested that *mdrL* presence did not significantly increase sanitizer tolerance, as it was present in all isolates, regardless of MIC. However, we cannot rule out whether differences in gene expression (62) may have been present in our isolates and contributed to differences in sanitizer tolerance.

The ListPred tool accurately predicted the low QAC tolerance of our isolates, which is unsurprising, as it is based on phenotypic QAC tolerance data from a large number of *L. monocytogenes* isolates primarily from Europe and the U.S. (39, 45, 63). It is less clear whether its virulence predictions are accurate (39). While the ListPred tool draws heavily on MLST-type clinical associations for virulence predictions, four of the five MLST types represented by our 30 isolates (STs 191, 361, 222, and 29) had fewer than 50 isolates in the Pasteur MLST database (32). While all had at least some clinical isolates-- 75% (3/4) of ST361 isolates were clinical, while 59.5% (22/37) of ST222 isolates, 55.8% (29/52) of ST451 isolates, and 22.2% of ST191 (2/9) isolates, respectively were clinical—the numbers were too small to draw any conclusions as to ListPred’s accuracy. Additionally, we did not find any mutations in key virulence genes, such as PMSCs in *inlA* (52, 64, 65), *inlB* deletions or truncations (66–68) or *prfA* mutations/truncations (69).

The capacity of *L. monocytogenes* to form either single-species and multi-species biofilms has been extensively assessed, but biofilm formation capacity is known to vary substantially among *L. monocytogenes* strains (4, 5, 14, 70). All 30 of the isolates tested had the capacity to attach to polystyrene plates, with no significant variations in the biomass of cell populations within the biofilms formed. Cell counts in the mature biofilms were not significantly different among isolates or between the ST191 persistent strain and the transient strains. However, in the SEM analysis of biofilm architecture, isolates FML-010 and FML-019 from the ST191/DUP-1042B persistent strains had substantially denser biofilms at 10 days, with strong 3D-development. Interestingly, we did not see the previously reported *L. monocytogenes* biofilm architecture of “knitted chains” or honeycombs (71, 72). Instead, we saw microcolonies with 3-D development, primarily found in surface micro blemishes for the weaker biofilm formers and the nutrient poor conditions. *L. monocytogenes* has been shown to preferentially inhabit surface blemishes and niches in processing environment materials, even more so than background microflora (73); and indeed, we primarily found dense biofilm clusters in surface micro blemishes.

Biofilm structure varied across the two nutrient conditions. For the strong biofilm formers (FML-009, FML-010, FML-029) in nutrient rich conditions, we typically saw dense, single-cell depth coverage, similar to the monospecies biofilms formed by environmental and clinical isolates in Rodriguez-Lopez et al.’s study (72);in Aryal et al. on glass slides (74); or the “strong” biofilm formers at two days growth at 30°C in Gamble and Muriana (75). Biofilms formed in nutrient poor conditions, however, had greater biomass, which included both cells and Extracellular Polymeric Substance, compared to nutrient-rich biofilms. Historic research indicates adherence can be increased in low nutrient conditions (15); however, we found that nutrient rich biofilms, especially when mature, had greater cell numbers. Given evidence from SEM images, we concluded that more EPS is produced in nutrient poor conditions than in nutrient rich conditions.

It was therefore not surprising that, despite the greater complexity observed in the biofilm architecture of nutrient rich biofilms, nutrient poor biofilms were equally resilient to sanitizer action.

Smaller cell numbers in nutrient poor biofilms were compensated for by greater EPS production which conferred protection against sanitizers. This is important in understanding how *L. monocytogenes* biofilms develop in nutrient poor food processing environments like non-food contact surfaces, examples of which include production floors, drains and cleaning equipment. It also highlights the importance of pre-sanitization steps like cleaning and abrasive action to effectively remove organic materials and EPS before application of sanitizers on non-food contact surfaces.

We did not find genetic mechanisms supporting our isolates tolerance to sanitizers either in planktonic or sessile forms. Given that QAC’s affect cell death by interfering with the cellular membrane (72, 76), resistance to the sanitizer in the biofilm samples may have been the product of protection conferred by the EPS. QACs are positively charged molecules which may have difficulty reaching the cells within the biofilms due to binding to the EPS, which is negatively charged (77). However, research comparing sanitizer tolerance in *Pseudomonas aeruginosa* and *Staphylococcus aureus* also suggests that *S. aureus*’ QAC susceptibility is reduced more in biofilms than *P. aeruginosa’s* (78), which suggests possible species-specific difference in biofilm susceptibility. Sodium hypochlorite, an active oxidizer, was significantly more effective against sessile cells than QACs. Nonetheless, it did not reach EPA-approved levels of 6 log reduction, which is required for claiming efficacy against biofilms (77). *L. monocytogenes* tolerance to SH has previously been shown to increase up to 100 times between early attachment (4 hours) and mature biofilms (79); consequently, it is vital to remove biofilms as quickly as possible.

A study in Austria investigated a persistent *L. monocytogenes* contamination scenario in a dairy processing plant (80) over a three-year period. They found that 19% of their samples were positive for *L. monocytogenes,* including samples from non-food contact surfaces, food contact surfaces, drain water, and shoes (80). As in the dairy our isolates came from, one subtype dominated the rest throughout the study, but other subtypes were mixed in (25, 80). Similarly, the frequency of *L. monocytogenes* isolation from shoes increased as time went on (80). Similarly, D’Amico and Donnelly only found *L. monocytogenes* on shoes in the final visit of their study (25, 26). In Austria, the same genotypes were found in the end products as on shoes (80). Ruckerl et al found no enhanced sanitizer tolerance in their study to QACs, hydrogen peroxide, or peracetic acid, though tolerance to QACs varied depending on the QAC (80). The authors did not assess biofilm formation capacity of their persistent isolates; however, they posited that biofilm formation likely contributed to the contamination of foodstuffs (80).

Another study investigated the transfer of *L. monocytogenes* from an older dairy processing facility to a new dairy processing facility, finding transfer of *L. monocytogenes* within just nine months of production beginning in the new facility (81). In this case, the authors found *L. monocytogenes* from a strain persistent in the first facility on the lids covering pallets of cheese for transfer to the second building (81). This persistent strain then became predominant in the second facility (81). Interestingly, the transferred persistent strain was not the major persistent strain in the first facility (81). However, its presence on food contact surfaces and in food facilitated its transfer into the new facility, where it rapidly colonized both food contact and non-food contact surfaces (81).

Both these studies describe processing plants with significant similarities to the one our ST191 isolates originated from: widespread, pervasive contamination, primarily on non-food contact surfaces, potentially uncontrolled traffic from the outside, and a dominant persistent strain in the facility that was difficult to eliminate (25, 26, 80, 81). It is crucial for dairy processers to establish traffic control SOPs to prevent *L. monocytogenes* colonization and to conduct frequent sampling and strain typing to detect colonizing strains and prevent them from establishing a firm foothold. A study in Brazil is instructive; the researchers identified *L. monocytogenes* in Minas Frescal Cheese, and traced the contamination back to the processing facility, where more than 40% of environmental samples were positive for *L. monocytogenes* (82). However, replacement of decrepit refrigerators and remodeling of the processing areas to reduce niches for *L. monocytogenes* successfully eliminated the pathogen for a very marginal cost to the producer (82). Consequently, early identification of contamination issues and prompt remediation may allow producers to avoid both outbreaks and prolonged shutdown periods for remediation.

## Conclusions

In the original studies by D’Amico and Donnelly, the authors posited that there might be enhanced sanitizer tolerance in the DUP-1042B isolates (25, 26); however, we did not observe this when we tested sanitizer tolerance. However, we discovered that the DUP-1042B isolates were developing enhanced attachment and biofilm formation capacity, which would have supported their persistence in the cheesemaking environment. We also identified more EPS in low-nutrient conditions. Consequently, non-food contact surfaces in cheese environments may induce the formation of biofilms with more EPS, which confers significant protection against sanitizers. The persistence of the ST-191/DUP-1042B strain of *L. monocytogenes* on non-food contact surfaces of raw milk artisanal cheese environments was therefore more likely the result of its enhanced biofilm formation than from an acquired genetic element.

## Acknowledgements

This research was conducted with funding from the George Walker Milk Fund, with additional salary support for Andrea Etter and Eurydice Aboagye from the Vermont Agricultural Experiment Station Hatch funding mechanism. We also thank and acknowledge the Vermont Department of Health for their assistance in sequencing these historic *L. monocytogenes* strains, Dr. Marc Allard of the FDA for his assistance in obtaining funding for the VDH through the Laboratory Flexible Funding Model (LFFM) Cooperative Agreement Program to sequence these environmental isolates, and Dr. Cathy Donnelly for transferring these strains to our custody upon her retirement.

---

**Figure S1:** graph of individual isolate MICs in sodium hypochlorite sanitizers (a) and QAC sanitizers (b) at nutrient rich and nutrient poor conditions

**Table S1:** List of isolates, with NCBI accession sequences.

## References

1. Hingston PA, Hansen LT, Pombert J-F, Wang S. Characterization of *Listeria monocytogenes* enhanced cold-tolerance variants isolated during prolonged cold storage. International Journal of Food Microbiology. 2019/10/02;306.

2. Cruz CD, Pitman AR, Harrow SA, Fletcher GC. *Listeria monocytogenes* associated with New Zealand seafood production and clinical cases: unique sequence types, truncated InlA, and attenuated invasiveness. Appl Environ Microbiol. 2014;80(4):1489–97.

3. Lunden JM, Autio TJ, Sjoberg AM, Korkeala HJ. Persistent and nonpersistent *Listeria monocytogenes* contamination in meat and poultry processing plants. J Food Prot. 2003;66(11):2062–9.

4. Vestby LK, Moretro T, Langsrud S, Heir E, Nesse LL. Biofilm forming abilities of *Salmonella* are correlated with persistence in fish meal- and feed factories. BMC Vet Res. 2009;5:20.

5. Wang J, Ray AJ, Hammons SR, Oliver HF. Persistent and transient *Listeria monocytogenes* strains from retail deli environments vary in their ability to adhere and form biofilms and rarely have *inlA* premature stop codons. Foodborne Pathog Dis. 2015;12(2):151–8.

6. Gray MJ, Zadoks RN, Fortes ED, Dogan B, Cai S, Chen Y, et al. *Listeria monocytogenes* isolates from foods and humans form distinct but overlapping populations. Appl Environ Microbiol. 2004;70(10):5833–41.

7. Gorski L, Cooley MB, Oryang D, Carychao D, Nguyen K, Luo Y, et al. Prevalence and clonal diversity of over 1,200 *Listeria monocytogenes* isolates collected from public access waters near produce production areas on the central California coast during 2011 to 2016. Appl Environ Microbiol. 2022;88(8):e0035722.

8. Scallan E, Hoekstra RM, Angulo FJ, Tauxe RV, Widdowson MA, Roy SL, et al. Foodborne illness acquired in the United States--major pathogens. Emerg Infect Dis. 2011;17(1):7–15.

9. Chen YI, Burall LS, Macarisin D, Pouillot R, Strain E, AJ DEJ, et al. Prevalence and level of *Listeria monocytogenes* in ice cream linked to a listeriosis outbreak in the United States. J Food Prot. 2016;79(11):1828–32.

10. Pouillot R, Klontz KC, Chen Y, Burall LS, Macarisin D, Doyle M, et al. Infectious dose of *Listeria monocytogenes* in outbreak linked to ice cream, United States, 2015. Emerg Infect Dis. 2016;22(12):2113–9.

11. Etter AJ, Hammons SR, Roof S, Simmons C, Wu T, Cook PW, et al. Enhanced sanitation standard operating procedures have limited impact on *Listeria monocytogenes* prevalence in retail delis. J Food Prot. 2017:1903–12.

12. Hammons SR, Etter AJ, Wang J, Wu T, Ford T, Howard MT, et al. Evaluation of third-party deep cleaning as a *Listeria monocytogenes* control strategy in retail delis. J Food Prot. 2017:1913–23.

13. Barria C, Singer RS, Bueno I, Estrada E, Rivera D, Ulloa S, et al. Tracing *Listeria monocytogenes* contamination in artisanal cheese to the processing environments in cheese producers in southern Chile. Food Microbiol. 2020;90:103499.

14. Nowak J, Cruz CD, Tempelaars M, Abee T, van Vliet AHM, Fletcher GC, et al. Persistent *Listeria monocytogenes* strains isolated from mussel production facilities form more biofilm but are not linked to specific genetic markers. Int J Food Microbiol. 2017;256:45–53.

15. Norwood DE, Gilmour A. Adherence of *Listeria monocytogenes* strains to stainless steel coupons. J Appl Microbiol. 1999;86(4):576–82.

16. Piercey MJ, Hingston PA, Truelstrup Hansen L. Genes involved in *Listeria monocytogenes* biofilm formation at a simulated food processing plant temperature of 15 degrees C. Int J Food Microbiol. 2016;223:63–74.

17. Ripolles-Avila C, Cervantes-Huaman BH, Hascoet AS, Yuste J, Rodriguez-Jerez JJ. Quantification of mature *Listeria monocytogenes* biofilm cells formed by an in vitro model: A comparison of different methods. Int J Food Microbiol. 2019;289:209–14.

18. Cherifi T, Arsenault J, Pagotto F, Quessy S, Cote JC, Neira K, et al. Distribution, diversity and persistence of *Listeria monocytogenes* in swine slaughterhouses and their association with food and human listeriosis strains. PLoS One. 2020;15(8):e0236807.

19. Nakari UM, Rantala L, Pihlajasaari A, Toikkanen S, Johansson T, Hellsten C, et al. Investigation of increased listeriosis revealed two fishery production plants with persistent *Listeria contamination* in Finland in 2010. Epidemiol Infect. 2014;142(11):2261–9.

20. CDC. Multistate outbreak of listeriosis linked to Blue Bell Creameries products (final update) 2015 [Available from: https://archive.cdc.gov/www_cdc_gov/listeria/outbreaks/ice-cream-03-15/index.html.

21. Flynn D. Vulto Creamery shut downbecause owner did not "understand". Food Safety News [Internet]. 2018. Available from: https://www.foodsafetynews.com/2018/04/vulto-creamery-shut-down-because-owner-did-not-understand/.

22. CDC. Multistate outbreak of listeriosis linked to Crave Brothers Farmstead Cheeses (final update) 2013 [Available from: https://archive.cdc.gov/www_cdc_gov/listeria/outbreaks/cheese-07-13/index.html.

23. Anonymous. Food Poisoning Bulletin [Internet]. Larsen L, editor: Eric Hageman. 2015 February 4, 2015. [cited 2024]. Available from: https://foodpoisoningbulletin.com/2015/cheese-listeria-outbreak-caused-2013-shutdown-of-crave-brothers/.

24. Paxson H. Cheese cultures: transforming American tastes and traditions. Gastronomica 2010;10(4):35–47.

25. D’Amico DJ, Donnelly CW. Enhanced detection of *Listeria spp*. in farmstead cheese processing environments through dual primary enrichment, PCR, and molecular subtyping. J Food Prot. 2008;71(11):2239–48.

26. D’Amico DJ, Donnelly CW. Detection, isolation, and incidence of *Listeria spp*. in small-scale artisan cheese processing facilities: a methods comparison. J Food Prot. 2009;72(12):2499–507.

27. Prjibelski A, Antipov D, Meleshko D, Lapidus A, Korobeynikov A. Using SPAdes *de novo* assembler. Curr Protoc Bioinformatics. 2020;70(1):e102.

28. Zerbino DR, Birney E. Velvet: algorithms for *de novo* short read assembly using de Bruijn graphs. Genome Res. 2008;18(5):821–9.

29. Zerbino DR. Using the Velvet *de novo* assembler for short-read sequencing technologies. Curr Protoc Bioinformatics. 2010;Chapter 11:Unit 11.5.

30. Wick RR, Judd LM, Gorrie CL, Holt KE. Completing bacterial genome assemblies with multiplex MinION sequencing. Microb Genom. 2017;3(10):e000132.

31. Larsen MV, Cosentino S, Rasmussen S, Friis C, Hasman H, Marvig RL, et al. Multilocus sequence typing of total-genome-sequenced bacteria. J Clin Microbiol. 2012;50(4):1355–61.

32. Anonymous. *Listeria* locus/sequence definitions: profiles. In: Pasteur I, editor. Institut Pasteur2024.

33. Carattoli A, Hasman H. PlasmidFinder and in Silico pMLST: identification and typing of plasmid replicons in whole-genome sequencing (WGS). Methods Mol Biol. 2020;2075:285–94.

34. Zankari E, Hasman H, Cosentino S, Vestergaard M, Rasmussen S, Lund O, et al. Identification of acquired antimicrobial resistance genes. J Antimicrob Chemother. 2012;67(11):2640–4.

35. Hanes RM, Huang Z. Investigation of antimicrobial resistance genes in *Listeria monocytogenes* from 2010 through to 2021. Int J Environ Res Public Health. 2022;19(9).

36. Joensen KG, Scheutz F, Lund O, Hasman H, Kaas RS, Nielsen EM, et al. Real-time whole-genome sequencing for routine typing, surveillance, and outbreak detection of verotoxigenic Escherichia coli. J Clin Microbiol. 2014;52(5):1501–10.

37. Expasy Translate Tool [Available from: https://web.expasy.org/translate/.

38. Gelbicova T, Kolackova I, Pantucek R, Karpiskova R. A novel mutation leading to a premature stop codon in inlA of *Listeria monocytogenes* isolated from neonatal listeriosis. New Microbiol. 2015;38(2):293–6.

39. Gmeiner A, Njage PMK, Hansen LT, Aarestrup FM, Leekitcharoenphon P. Predicting *Listeria monocytogenes* virulence potential using whole genome sequencing and machine learning. Int J Food Microbiol. 2024;410:110491.

40. Hallgren MB, Overballe-Petersen S, Lund O, Hasman H, Clausen P. MINTyper: an outbreak-detection method for accurate and rapid SNP typing of clonal clusters with noisy long reads. Biol Methods Protoc. 2021;6(1):bpab008.

41. Muchaamba F, Guldimann C, Tasara T, Mota MI, Braga V, Varela G, et al. Full-genome sequence of *Listeria monocytogenes* strain H34, isolated from a newborn with sepsis in Uruguay. Genome Announc. 2017;5(24).

42. Kaas RS, Leekitcharoenphon P, Aarestrup FM, Lund O. Solving the problem of comparing whole bacterial genomes across different sequencing platforms. PLoS One. 2014;9(8):e104984.

43. Ryan S, Begley M, Hill C, Gahan CG. A five-gene stress survival islet (SSI-1) that contributes to the growth of *Listeria monocytogenes* in suboptimal conditions. J Appl Microbiol. 2010;109(3):984–95.

44. Bankevich A, Nurk S, Antipov D, Gurevich AA, Dvorkin M, Kulikov AS, et al. SPAdes: a new genome assembly algorithm and its applications to single-cell sequencing J Comput Biol. 2012;19(5):455.

45. Ivanova M, Kragh ML, Szarvas J, Tosun E, Holmud N, Gmeiner A, et al. Large-scale phenotypic and genomic analysis of *Listeria monocytogenes* reveals diversity in the sensitivity to quaternary ammonium compounds but not to peracetic acid. BioRxiv Preprint. 2023.

46. Borucki MK, Kim SH, Call DR, Smole SC, Pagotto F. Selective discrimination of *Listeria monocytogenes* epidemic strains by a mixed-genome DNA microarray compared to discrimination by pulsed-field gel electrophoresis, ribotyping, and multilocus sequence typing. J Clin Microbiol. 2004;42(11):5270–6.

47. Jagadeesan B, Baert L, Wiedmann M, Orsi RH. Comparative analysis of tools and approaches for source tracking *Listeria monocytogenes* in a food facility using whole-genome sequence data. Front Microbiol. 2019;10:947.

48. Orsi RH, Borowsky ML, Lauer P, Young SK, Nusbaum C, Galagan JE, et al. Short-term genome evolution of *Listeria monocytogene*s in a non-controlled environment. BMC Genomics. 2008;9:539.

49. Stasiewicz MJ, Oliver HF, Wiedmann M, den Bakker HC. Whole-genome sequencing allows for improved identification of persistent *Listeria monocytogenes* in food-associated environments. Appl Environ Microbiol. 2015;81(17):6024–37.

50. Holch A, Webb K, Lukjancenko O, Ussery D, Rosenthal BM, Gram L. Genome sequencing identifies two nearly unchanged strains of persistent *Listeria monocytogenes* isolated at two different fish processing plants sampled 6 years apart. Appl Environ Microbiol. 2013;79(9):2944–51.

51. Pasquali F, Palma F, Guillier L, Lucchi A, De Cesare A, Manfreda G. *Listeria monocytogenes* sequence types 121 and 14 repeatedly isolated within one year of sampling in a rabbit meat processing plant: persistence and ecophysiology. Front Microbiol. 2018;9:596.

52. Moura A, Criscuolo A, Pouseele H, Maury MM, Leclercq A, Tarr C, et al. Whole genome-based population biology and epidemiological surveillance of *Listeria monocytogenes*. Nat Microbiol. 2016;2:16185.

53. Anonymous. In: Pasteur I, editor.

54. Kim SW, Haendiges J, Keller EN, Myers R, Kim A, Lombard JE, et al. Genetic diversity and virulence profiles of *Listeria monocytogenes* recovered from bulk tank milk, milk filters, and milking equipment from dairies in the United States (2002 to 2014). PLoS One. 2018;13(5):e0197053.

55. Cardenas-Alvarez MX, Zeng H, Webb BT, Mani R, Munoz M, Bergholz TM. Comparative genomics of *Listeria monocytogenes* isolates from ruminant listeriosis cases in the midwest United States. Microbiol Spectr. 2022;10(6):e0157922.

56. Steckler AJ, Cardenas-Alvarez MX, Townsend Ramsett MK, Dyer N, Bergholz TM. Genetic characterization of *Listeria monocytogenes* from ruminant listeriosis from different geographical regions in the U.S. Vet Microbiol. 2018;215:93–7.

57. Felix B, Capitaine K, Te S, Felten A, Gillot G, Feurer C, et al. Identification by high-throughput real-time PCR of 30 major circulating *Listeria monocytogenes* clonal complexes in Europe. Microbiol Spectr. 2023;11(3):e0395422.

58. Kurpas M, Osek J, Moura A, Leclercq A, Lecuit M, Wieczorek K. Genomic characterization of *Listeria monocytogenes* isolated from ready-to-eat meat and meat processing environments in Poland. Front Microbiol. 2020;11:1412.

59. Brauge T, Leleu G, Hanin A, Capitaine K, Felix B, Midelet G. Genetic population structure of *Listeria monocytogenes* strains isolated from salmon and trout sectors in France. Heliyon. 2023;9(7):e18154.

60. Ricchi M, Scaltriti E, Cammi G, Garbarino C, Arrigoni N, Morganti M, et al. Short communication: Persistent contamination by *Listeria monocytogenes* of bovine raw milk investigated by whole-genome sequencing. J Dairy Sci. 2019;102(7):6032–6.

61. Jiang X, Yu T, Xu Y, Wang H, Korkeala H, Shi L. MdrL, a major facilitator superfamily efflux pump of *Listeria monocytogenes* involved in tolerance to benzalkonium chloride. Appl Microbiol Biotechnol. 2019;103(3):1339–50.

62. Romanova NA, Wolffs PF, Brovko LY, Griffiths MW. Role of efflux pumps in adaptation and resistance of *Listeria monocytogenes* to benzalkonium chloride. Appl Environ Microbiol. 2006;72(5):3498–503.

63. Gmeiner A, Ivanova M, Njage PMK, Hansen LT, Chindelevitch L, Leekitcharoenphon P. Quantitative prediction of disinfection tolerance in *Listeria monocytogenes* using whole genome sequencing and machine learning. BioRxiv Preprint. 2023.

64. Jacquet C, Doumith M, Gordon JI, Martin PM, Cossart P, Lecuit M. A molecular marker for evaluating the pathogenic potential of foodborne *Listeria monocytogenes*. J Infect Dis. 2004;189(11):2094–100.

65. Maury MM, Tsai YH, Charlier C, Touchon M, Chenal-Francisque V, Leclercq A, et al. Uncovering *Listeria monocytogenes* hypervirulence by harnessing its biodiversity. Nat Genet. 2016;48(3):308–13.

66. Liu D, Bai X, Helmick HDB, Samaddar M, Amalaradjou MAR, Li X, et al. Cell-surface anchoring of *Listeria* adhesion protein on *L. monocytogenes* is fastened by *internalin B* for pathogenesis. Cell Rep. 2023;42(5):112515.

67. Zakrzewski AJ, Chajecka-Wierzchowska W, Zadernowska A, Podlasz P. Virulence characterization of *Listeria monocytogenes, Listeria innocua*, and *Listeria welshimeri* isolated from fish and shrimp using *in vivo* early zebrafish larvae models and molecular study. Pathogens. 2020;9(12).

68. Sobyanin K, Sysolyatina E, Krivozubov M, Chalenko Y, Karyagina A, Ermolaeva S. Naturally occurring InlB variants that support intragastric *Listeria monocytogenes* infection in mice. FEMS Microbiol Lett. 2017;364(3).

69. Roche SM, Gracieux P, Milohanic E, Albert I, Virlogeux-Payant I, Temoin S, et al. Investigation of specific substitutions in virulence genes characterizing phenotypic groups of low-virulence field strains of *Listeria monocytogenes*. Appl Environ Microbiol. 2005;71(10):6039–48.

70. Harvey J, Keenan KP, Gilmour A. Assessing biofilm formation by *Listeria monocytogenes* strains. Food Microbiol. 2007;24(4):380–92.

71. Rieu A, Briandet R, Habimana O, Garmyn D, Guzzo J, Piveteau P. *Listeria monocytogenes* EGD-e biofilms: no mushrooms but a network of knitted chains. Appl Environ Microbiol. 2008;74(14):4491–7.

72. Rodriguez-Lopez P, Rodriguez-Herrera JJ, Lopez Cabo M. Architectural features and resistance to food-grade disinfectants in *Listeria monocytogenes-Pseudomonas* spp. dual-species biofilms. Front Microbiol. 2022;13:917964.

73. Fagerlund A, Moretro T, Heir E, Briandet R, Langsrud S. Cleaning and disinfection of biofilms composed of *Listeria monocytogenes* and background microbiota from meat processing surfaces. Appl Environ Microbiol. 2017;83(17).

74. Aryal M, Muriana PM. Efficacy of commercial sanitizers used in food processing facilities for inactivation of *Listeria monocytogenes, E. coli* O157:H7, and *Salmonella* biofilms. Foods. 2019;8(12).

75. Gamble R, Muriana PM. Microplate fluorescence assay for measurement of the ability of strains of *Listeria monocytogenes* from meat and meat-processing plants to adhere to abiotic surfaces. Appl Environ Microbiol. 2007;73(16):5235–44.

76. Perez-Rodriguez M, Lopez Cabo M, Balsa-Canto E, Garcia MR. Mechanisms of *Listeria monocytogenes* disinfection with benzalkonium chloride: from molecular dynamics to kinetics of time-kill curves. Int J Mol Sci. 2023;24(15).

77. Lineback CB, Nkemngong CA, Wu ST, Li X, Teska PJ, Oliver HF. Hydrogen peroxide and sodium hypochlorite disinfectants are more effective against *Staphylococcus aureus* and *Pseudomonas aeruginosa* biofilms than quaternary ammonium compounds. Antimicrob Resist Infect Control. 2018;7:154.

78. Campanac C, Pineau L, Payard A, Baziard-Mouysset G, Roques C. Interactions between biocide cationic agents and bacterial biofilms. Antimicrob Agents Chemother. 2002;46(5):1469–74.

79. Lee SH, Frank JF. Inactivation of surface-adherent *Listeria monocytogenes* hypochlorite and heat. J Food Prot. 1991;54(1):4–6.

80. Ruckerl I, Muhterem-Uyar M, Muri-Klinger S, Wagner KH, Wagner M, Stessl B. *L. monocytogenes* in a cheese processing facility: learning from contamination scenarios over three years of sampling. Int J Food Microbiol. 2014;189:98–105.

81. Melero B, Stessl B, Manso B, Wagner M, Esteban-Carbonero OJ, Hernandez M, et al. *Listeria monocytogenes* colonization in a newly established dairy processing facility. Int J Food Microbiol. 2019;289:64–71.

82. Brito JR, Santos EM, Arcuri EF, Lange CC, Brito MA, Souza GN, et al. Retail survey of Brazilian milk and Minas frescal cheese and a contaminated dairy plant to establish prevalence, relatedness, and sources of *Listeria monocytogenes* isolates. Appl Environ Microbiol. 2008;74(15):4954–61.

83. Keeney K, Trmcic A, Zhu Z, Delaquis P, Wang S. Stress survival islet 1 contributes to serotype-specific differences in biofilm formation in *Listeria monocytogenes*. Lett Appl Microbiol. 2018;67(6):530–6.

